# Assessment of research ethics education offerings of pharmacy master programs: a qualitative content analysis

**DOI:** 10.1101/2020.08.25.266023

**Authors:** Wesam S. Ahmed, Camille Nebeker

## Abstract

The importance of research ethics (RE) training has led academic and funding institutions to require that students, trainees, and faculty obtain such training at various stages of their careers. Despite the increasing awareness of the value RE education offers, this training requirement is absent in Jordan. We aimed to assess RE education offerings of pharmacy master programs in Jordan and compare with the top-ranked pharmacy graduate programs globally. Therefore, a list of universities that offer research-based pharmacy master programs was created. Each program was evaluated for the inclusion of RE education. A qualitative content analysis approach based on inductive reasoning and latent analysis was followed to analyze the data. Results of the study showed a lack of appropriate RE education for graduate-level pharmacy programs in Jordan with only 40% of the programs partially discuss selected topics related to RE. Regarding pharmacy graduate programs globally, 10% offer a standalone RE course, 40% offer some discussions related to RE, another 10% do not offer RE education in any form, and the remaining 40% of the programs were difficult to assess due to lack of sufficient information available online. Based on the findings of this study, training in RE is tends to be lacking in pharmacy graduate programs in Jordan and globally, with a greater lack in Jordan than globally. There is a need to incorporate formal RE education into programs that do not offer this type of instruction. Programs that formally touch on some aspects of RE need to expand the scope of topics to include more RE-related themes. Integrating a standalone RE course into pharmacy graduate programs is highly encouraged.

## Introduction

The United States National Institute of Health (NIH) requirement for education in the responsible conduct of research (RCR) [1] states that the practice of science with integrity involves “*the awareness and application of established professional norms and ethical principles in the performance of all activities related to scientific research*”. Clearly, ethical behavior in science is valued, and it would follow that research ethics education is the modality for acculturating scientists to the accepted norms and conventions. In 1989, the NIH announced its first RCR educational requirement for selected NIH trainees [2]. Although increased cases of research misconduct may have triggered the federal requirements for RCR training [3, 4], the desire to preserve the integrity of science and to foster a good research practices and a socially responsible community could arguably be another reason [5–9]. The educational requirements were expanded in 1992 and 2000 for the NIH [10, 11] and by the Public Health Service (PHS) in 2000 (http://grants.nih.gov/grants/guide/notice-files/NOT-OD-01-007.html). The NIH updated its mandate for RCR training in 2009 [1] and, that same year, the National Science Foundation (NSF) introduced an RCR requirement that called for “appropriate training and oversight in the responsible and ethical conduct of research” [12]. In 2010, the NSF RCR requirement went into effect, which required that institutions receiving NSF support have a plan for offering RCR education for all students/trainees (undergraduate, graduate, and postdoctoral) supported on NSF grants. Recently, the National Institute of Food and Agriculture (NIFA) of the United States Department of Agriculture (USDA) incorporated RCR education as an essential requirement for institution conducting USDA-funded extramural research (https://nifa.usda.gov/responsible-andethical-conduct-research).

Unlike the NSF requirement, which only calls for “appropriate training” to be provided to researchers supported by the NSF, the NIH requirement includes detail on what topics to be discussed, as well as expectations on format and frequency of training [4]. The scope of RCR training requirements has varied over different versions of NIH mandates (1989, 1992, 2000, and 2009). In fact, the U.S. office of research integrity (ORI) identified nine core areas which need to be addressed in an RCR course. This was followed by a Delphi consensus panel report which identified 53 topics in seven core areas to be included in RE teaching [13]. Some RCR topics have evolved over time, while others were newly introduced in later versions of the federal requirements. Generally, accepted topics include conflict of interest and bias, research subject protections, data management, authorship and publication ethics, and social responsibilities [4].

As noted, increased cases of research misconduct in the 1980’s led to a special congressional task force to define the scope of research misconduct [3, 4] and, eventually, federal requirements for RCR training. How the training requirements were implemented by institutions bound by these new training requirements varied considerably across the US [13–15]. In addition to the variability in RCR instruction by institution, inconsistencies in what instructors thought they were to accomplish specific to goals were also common [16] making evaluation of the efficacy of RCR training challenging [5]. A range of suggested goals are available in the RCR literature [13, 16].

While the federal agencies mandate institutions to provide RCR training to comply with training requirements; increasingly, institutions offer RCR training regardless of funding requirements [4]. In addition to U.S. institutions promoting research ethics training to foster an ethically responsible research environment, the NIH Fogarty International Center (FIC) also supports proactive research ethics education internationally. For example, FIC has funded research ethics education in Latin America, Africa and in the Middle East. Current research practices in most Arab countries are not governed by nation-level federal requirements nor do they involve the same level of mandates for research ethics as the U.S. [17]. Seeing as many collaboration research projects in these countries are funded by U.S. agencies [18], this could be a reasonable motive behind FIC funds outside of the U.S.

The situation in Jordan is different from the U.S., yet similar to most Arab countries in that there are no research ethics education requirements for students and researchers regardless of research funding source. Nonetheless, Jordan is considered one of the most academically established countries in the MENA region with progressive research agendas. Jordan also has a well-established pharmaceutical industry that exports several products globally. This pharmaceutical sector relies heavily on contract research organizations for drug development and post-marketing studies [17]. As a result, Jordan has been part of two major initiatives of research ethics training, both of which are supported by grants from the NIH FIC:

- Middle East Research Ethics Training Initiative (MERETI), initiated in 2005, and offers training in research ethics to individuals (mid-level and senior professionals) from the Middle East. (https://www.mereti-network.net/; https://www.fic.nih.gov/Grants/Search/Pages/Bioethics-1R25TW007090-01.aspx)
- The Research ethics education program in Jordan (REPJ), which started in 2015, and targets junior researchers from the MENA region. (https://jordanrcrprogram.com/about/; https://www.fic.nih.gov/Grants/Search/Pages/bioethics-TW010026.aspx)

Graduate masters-level pharmacy programs in Jordan generally require that students complete an independent research project. Given this expectation and the potential value of RCR education in preparing researchers to design and implement their research ethically and responsibly, we sought to assess the extent to which research ethics education was included in graduate pharmacy programs in Jordan. To see how Jordan pharmacy programs compared globally with respect to RCR offerings, we then sought investigate the offerings at the top ranked graduate pharmacy programs.

## Materials and methods

### Data collection in Jordan

Data collection for this part involved several steps. In the begining, websites of all Jordanian universities with a pharmacy school or department were reviewed to identify schools/departments offering one or more masters-level programs in pharmacy. Then, we searched through each program’s description to document programs requiring students to carry out an independent research project to satisfy completion requirements along with the course-work requirement. In case the program’s description was not available online, it was retrieved from the corresponding pharmacy school/department by a direct in-person visit. Next, we searched through the description of individual courses in a program to identify courses that offer research ethics education. The purpose was to identify whether programs that included an expectation to conduct research also provided RE instruction. Based on this search, a spreadsheet was compiled (S1 Table) documenting courses with dedicated or imbedded material related to research ethics by examining the list of course titles and their descriptions. Research ethics instruction was determined by searching for keywords related to research ethics in the course description. For courses that contained one or more keywords related to research ethics in their description, in order to determine the scope of research ethics topics covered, the entire course content was then retrieved by a direct in-person visit to the corresponding pharmacy school/department. Fig 1 summarizes the data collection process in Jordan.

**Fig 1.**
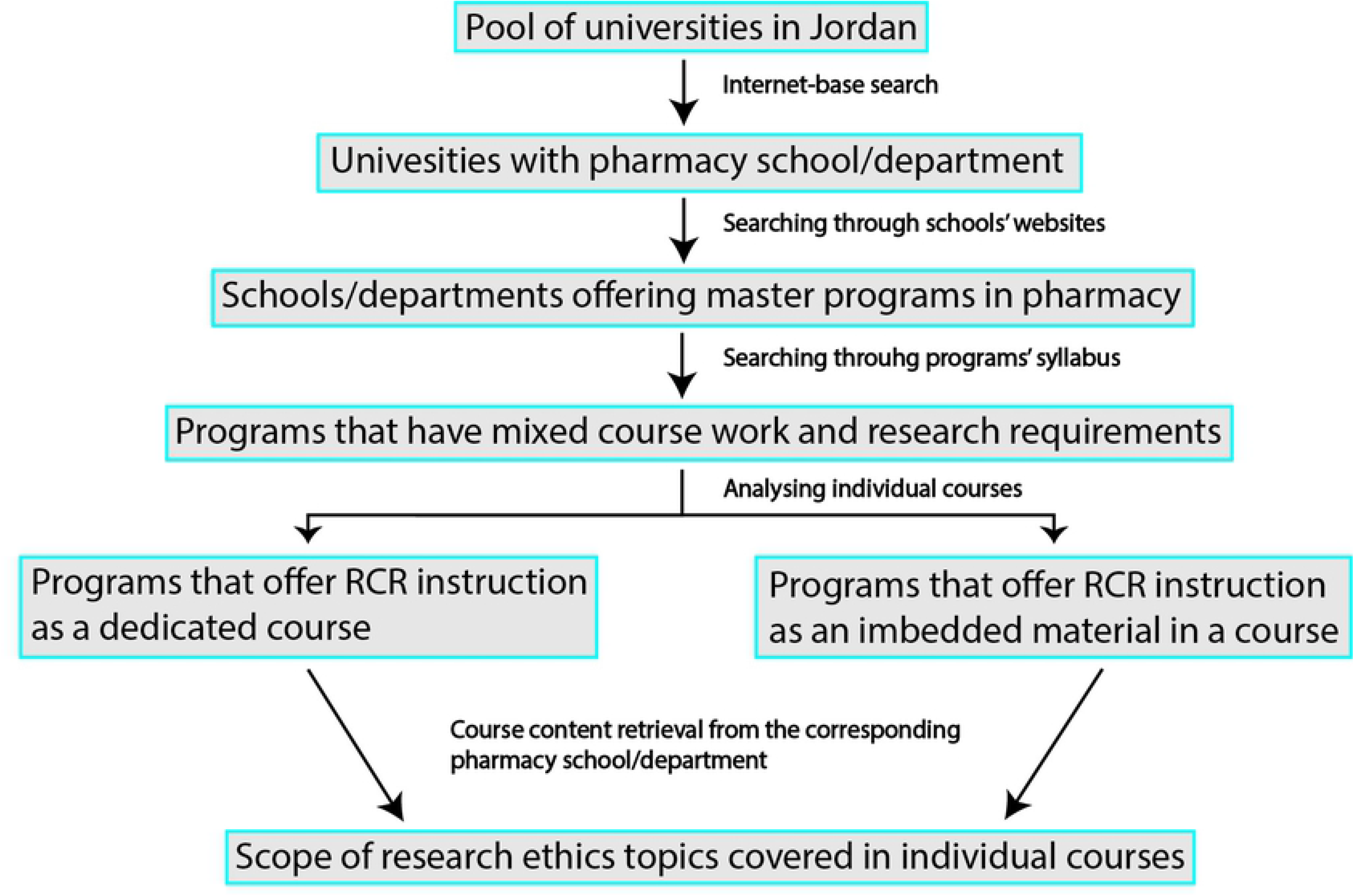
Stepwise data collection approach from universities in Jordan.

Data collected during the review process were entered into a spreadsheet included: 1- name of the university and the master pharmacy program offered, 2- name of all courses; core (obligatory) and elective (optional) offering research ethics instruction, 3- course description, 4- keywords related to research ethics used in the course description, 5- whether research ethics are the main and the only focus of the course or an imbedded material (i.e. research ethics are mentioned or discussed in the course but are not the only focus of the course), and 6- research ethics related topics covered in the course(S1 Table).

### Data collection beyond Jordan

The next step was to compare graduate research ethics educational offerings in Jordan with that of pharmacy graduate programs offered globally. For this part, the data collection was based entirely on information available online. Since our data collection took place during July – August 2017, we therefore used the Quacquarelli Symonds (QS) World University Ranking (2017) to search for the top 10 universities worldwide by subject of pharmacy and pharmacology (https://www.topuniversities.com/university-rankings/university-subject-rankings/2017/pharmacy-pharmacology). The QS World University Ranking, published by Quacquarelli Symonds (QS) Limited, is one of the most popular and reputable rankings in the educational market [19].

After the list of the top 10 universities was created, websites of these universities were then reviewed for pharmacy master programs that have mixed coursework and research-project completion requirements. Individual programs were then screened by reviewing the course description of individual courses in the programs for core or elective courses that fully or partly discuss research ethics related issues. The data collection process of global programs is summarized in Fig 2.

**Fig 2.**
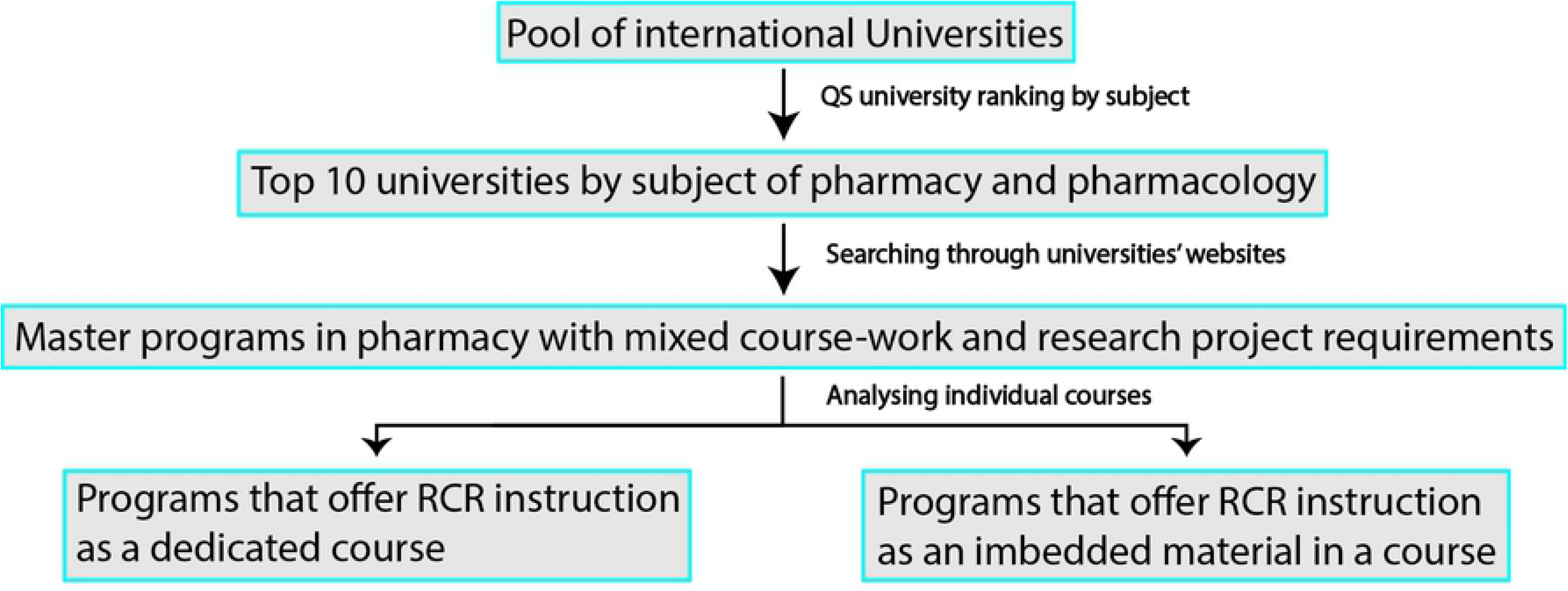
Stepwise data collection approach from top 10 universities globally.

Based on this search, data that were collected and recorded in a spreadsheet and included: 1- name and rank of the university, 2- name of the master program offered, 3- the school or department offering the program, 4- course title, 5- whether the course is core or elective, 6- the course description, 7- keywords related to research ethics used in the course description, 8-whether the course offers research ethics instruction as the only focus or as one component of the course, and 9- the course website address (the website address of the program was used whenever no specific website address is available for the course) [S2 Table]. All information was retrieved from official online sources with no direct or indirect contact with the universities or their schools.

### Data analysis

The study aim was to qualify and quantify research ethics instruction offered by pharmacy master programs in Jordan and compare the results with the top 10 universities by subject of pharmacy and pharmacology. Our first sample included pharmacy master programs with mixed coursework and research completion requirements that are offered by universities in Jordan. Our second sample included master programs offered by the top 10 universities by subject of pharmacy and pharmacology. Individual courses in the programs served as units of analysis, courses discerptions served as meaning units, and keywords observations from written course description texts served as condensed meaning units. A qualitative content analysis, based on inductive reasoning was applied [20]. Several other studies utilized content analysis to assess major text books and other educational resources for RCR content [21, 22]. We followed a latent analysis approach, as some of the keywords we extracted from the description texts to reflect research ethics instruction may not explicitly refer to research ethics in an obvious manner, rather it was our interpretation of these keywords through which we tried to seek their underlying content reference.

In order to conceptualize on the collected data, the data analysis process consisted of the following four steps: the decontextualization, the recontextualization, the categorization and theming, and the compilation (Fig 3). The decontextualization step entails reading though texts to identify meaning units, which are broken down into condensed meaning units, and creating a coding list. Recontextualization includes comparing all meaning units from the previous step with the original text to check and make sure that all aspects of the relevant content have been captured and coded properly. Categorization involves grouping the created codes into subcategories and categories that are homogenous on the interior but heterogenous on the exterior, that are then appropriately themed. The compilation is the last step through which the results are compiled into meaningful conclusions.

**Fig 3.**
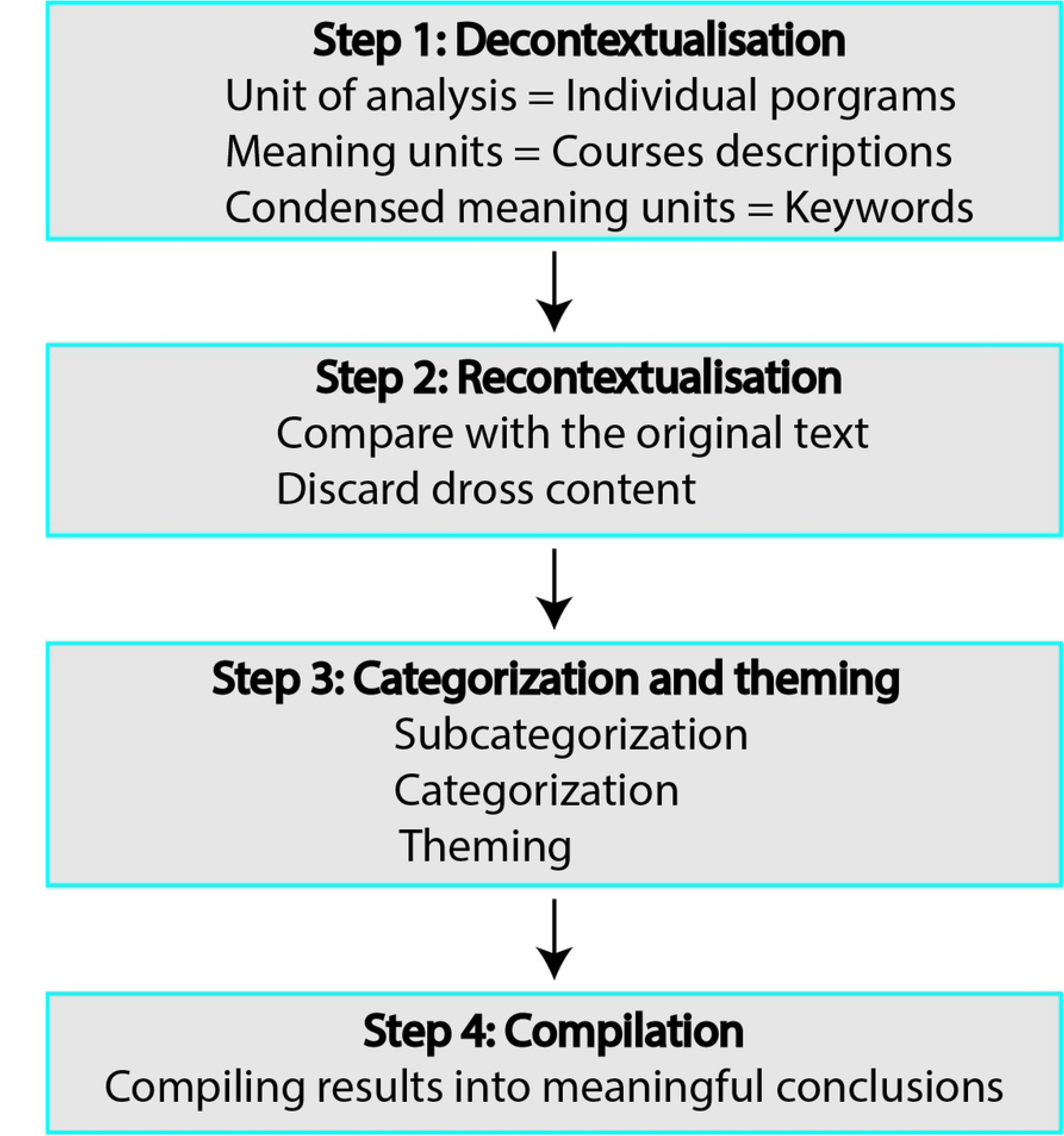
Overview of the data analysis process.

Step 1. Decontextualisation: For each course, the course description served as the meaning unit, keywords related to research ethics served as condensed meaning unit. A coding list was created for each course as follows:

I. If keywords were observed in the description and the course title was indicative of a course specialized in research ethics (e.g. “research ethics” course, “responsible conduct of research” course), we coded “Yes - dedicated” for that course.
II. If the description included keywords related to research ethics, but the course title and description were indicative of other contents unrelated to research ethics, we coded “Yes – imbedded” for that course.
III. If the course did not include any keywords related to research ethics, we coded “No” for that course.

Step 2. Recontextualisation: The condensed meaning units were put back and compared with the original description text to make sure all relevant keywords related to research ethics have been captured. Then, all other text words in the description were considered dross and excluded from further analysis.

Step 3. Categorization and theming:

I. Subcategories

a. Core and elective courses coded as “Yes – dedicated”.
b. Core and elective courses coded as “Yes – imbedded”.
c. Core and elective courses coded as “No”.
II. Categories

a. Programs that included subcategory (a) or (b).
b. Programs that included subcategory (a).
c. Programs that included subcategory (b) but not (a).
d. Programs that included subcategory (c) but not (a) or (b).
e. Programs that did not include any of the subcategories (a), (b), or (c).
III. Themes

Category (a) was themed as programs that offer some form of research ethics education in one or more of their courses.
Category (b) was themed as programs that offer one or more dedicated research ethics course.
Category (c) was themed as programs that offer research ethics education imbedded into one or more of their courses.
Category (d) was themed as programs that do not offer research ethics education in any form.
Category (e) was themed as programs that were difficult to assess for research ethics education offerings due to lack of sufficient information online.

Step 4. Compilation: results were compiled and used to draw meaningful conclusions as discussed later in this article.

## Results

### RCR in Jordan

Our search revealed 19 universities in Jordan that have a pharmacy school or department with only 7 of those offering mixed course-work and research-project master’s degree in pharmacy (Table 2). The total number of pharmacy master programs offered in the 7 universities was 10 (Table 3). All 10 programs stated a completion requirement to conduct a research project. With the exception of the University of Jordan (JU) and Jordan University of Science and Technology (JUST), all other universities that offer pharmacy master’s degree programs, reported offering neither a dedicated course on research ethics nor research ethics-integrated formal discussions within an existing course. The school of pharmacy at the University of Jordan (JU) offers a Master of Science (MSc) Clinical Pharmacy program with one obligatory course in “research methodology” that includes RE instruction as one component of the course. The imbedded RE material is focused on research involving humans with emphasis on informed consent and role of an institutional review board, authorship and publication ethics, conflict of interest, data confidentiality, and research misconduct (fabrication, falsification, and plagiarism) [S1 Table]. Most of these topics align with the NIH federal requirements for RCR. The master program in Pharmaceutical Sciences offered by the same school, however, does not discuss research ethics in any of its courses, despite having a research project completion mandate. The school of pharmacy at Jordan University of Science and Technology (JUST) offers three master programs, each with an obligatory “research methodology” course, which includes some discussion of ethical issues related to research. The same topics were covered in the three courses with focus on human research, animal research, and research misconduct, all of which align with the NIH mandates (Table 3 and S1 Table).

**Table 2.**
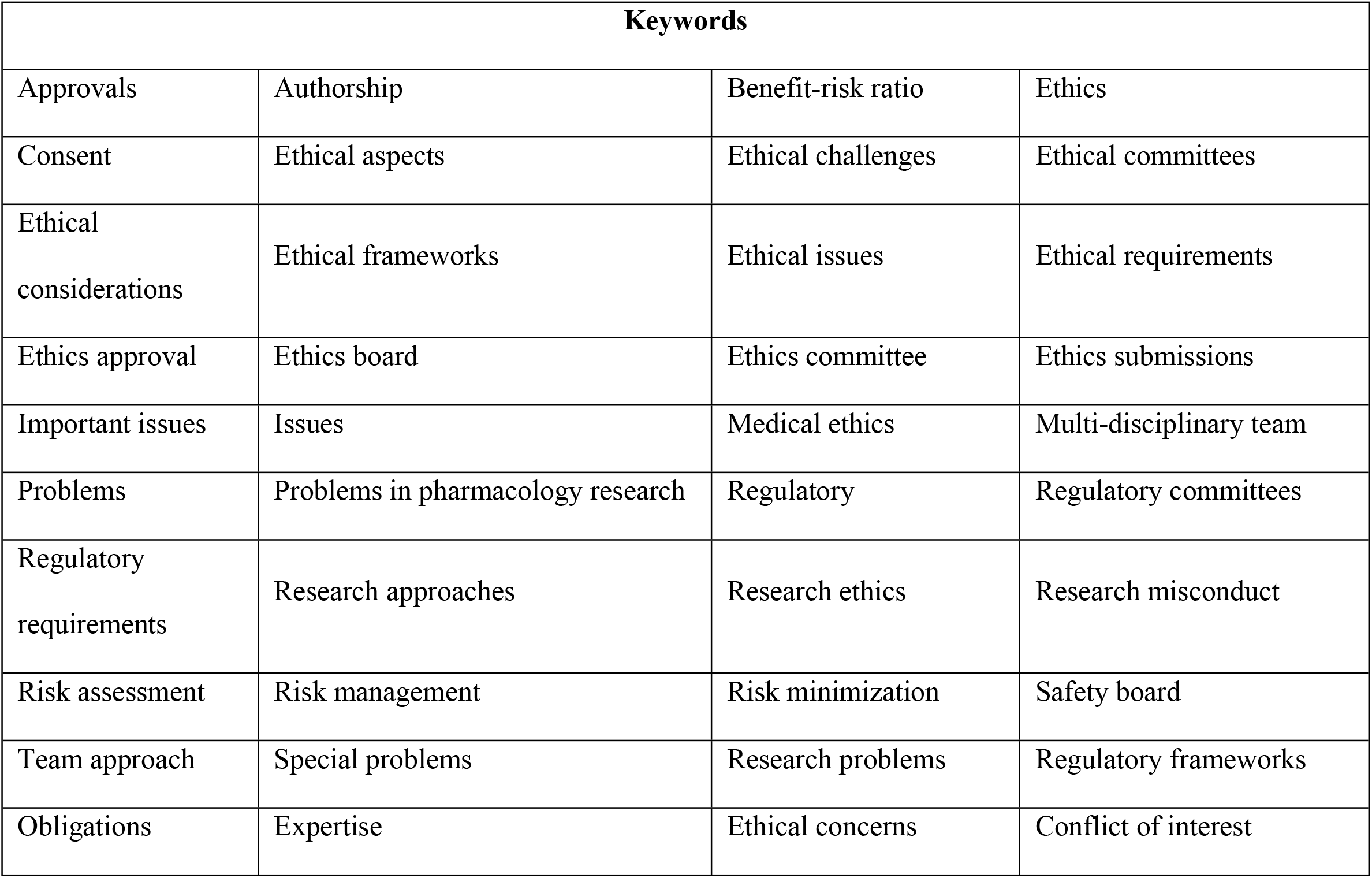
Keywords related to research ethics that were highlighted from course discerptions.

**Table 2.** List of Universities with Pharmacy Schools/Departments in Jordan.

| # | Name of University | Private University | City | Pharmacy master program(s) with thesis track |
| --- | --- | --- | --- | --- |
| 1 | Jordan University of Science and Technology | No | Irbid | I. Clinical Pharmacy<br>II. Pharmaceutical Technology<br>III. Medicinal chemistry and Pharmacognosy |
| 2 | University of Jordan | No | Amman | I. Clinical Pharmacy<br>II. Pharmaceutical Sciences |
| 3 | Yarmouk University | No | Irbid | None |
| 4 | Hashemite University | No | Zarqa | None |
| 5 | Zarqa University | Yes | Zarqa | None |
| 6 | Balqa Applied University |  | Irbid | None |
| 7 | Philadelphia University at Jordan | Yes | Jarash | None |
| 8 | Mutah University | No | Karak | None |
| 9 | German Jordanian University | No | Amman | None |
| 10 | Applied Science University | Yes | Amman | Pharmaceutical Sciences |
| 11 | Al Ahliyya Amman University | Yes | Balqa | Pharmaceutical Studies |
| 12 | Middle East University Jordan | Yes | Amman | None |
| 13 | Al Zaytoonah University | Yes | Amman | Pharmaceutical Sciences |
| 14 | University of Petra | Yes | Amman | Pharmaceutical Sciences |
| 15 | Jerash Private University | Yes | Jarash | None |
| 16 | Amman Arab University | Yes | Amman | None |
| 17 | American University of Madaba | Yes | Madaba | None |
| 18 | Al Isra University Amman | Yes | Amman | Pharmacy |
| 19 | Jadara University | Yes | Irbid | None |

**Table 3.** Pharmacy master programs by mixed course-work/research offered by Jordanian universities and their research ethics (RE) education offerings.

| Pharmacy master program | RE courses offered |
| --- | --- |

|  | No | Yes |  | N/A |
| --- | --- | --- | --- | --- |
|  |  | Dedicated (C, E) | Integrated (C, E) |  |
| *The University of Jordan* |  |  |  |  |
| Clinical Pharmacy | - | 0 | 1 (C) | - |
| Pharmaceutical Sciences | ✓ | 0 | 0 | - |
| *Jordan university of Science and Technology* |  |  |  |  |
| Clinical Pharmacy | - | 0 | 1 (C) | - |
| Pharmaceutical Technology | - | 0 | 1 (C) | - |
| Medicinal Chemistry and pharmacognosy | - | 0 | 1 (C) | - |
| *Applied Science University* |  |  |  |  |
| Pharmaceutical Studies | ✓ | 0 | 0 |  |
| *Al Ahliyya Amman University* |  |  |  |  |
| Pharmaceutical Studies | ✓ | 0 | 0 | - |
| *Al Zaytoonah University* |  |  |  |  |
| Pharmaceutical Sciences | ✓ | 0 | 0 | - |
| *University of Petra* |  |  |  |  |
| Pharmaceutical Sciences | ✓ | 0 | 0 | - |
| *Al Isra University Amman* |  |  |  |  |
| Pharmacy | ✓ | 0 | 0 | - |

Dedicated: number of standalone RE courses offered by a program; Integrated: number of courses in a program that integrate RE discussions; N/A: no enough information to assess RE education offerings; C: Core course; E: elective course.

### RCR beyond Jordan - globally

According to the information available online, the total number of pharmacy master programs with mixed course-load and research requirement offered by the top 10 universities by subject of pharmacy and pharmacology is 20 programs. Of the 20 programs, two programs (10%) offer a dedicated research ethics course, eight programs (40%) offer research ethics education material that is integrated into one or more of the program’s courses, and two programs (10%) contain neither a dedicated course of research ethics nor a course that integrates research ethics material. Another eight programs (40%) were difficult to assess for research ethics instruction offerings because the course description of these programs was either missing or incomplete. (Table 4 and S2 Table).

**Table 4.** Pharmacy master programs by mixed course-work/research offered by the top 10 universities ranked by subject of pharmacy and pharmacology (QS ranking, 2017) and their research ethics (RE) education offerings. Universities are listed based on their ranking in the top 10 list (highest to lowest).

| Pharmacy master program | RE courses offered |  |  |  |
| --- | --- | --- | --- | --- |
|  | No | Yes |  | N/A |
|  |  | Dedicated (C, E) | Integrated (C, E) |  |
| *University of Harvard (no programs are offered)* |  |  |  |  |
| *University of Monash* |  |  |  |  |
| Master of Clinical Pharmacy | - | 0 | 1 (C), 1 (E) | - |
| *University of Cambridge (no mixed programs are offered)* |  |  |  |  |
| *University of Oxford* |  |  |  |  |
| Pharmacology | - | - | - | ✓ |
| *University of California, San Francisco* |  |  |  |  |
| Clinical Research | - | 1 (C) | 0 | - |
| *University of Nottingham* |  |  |  |  |
| Drug Discovery and Pharmaceutical Sciences | - | - | - | ✓ |
| *King's College London* |  |  |  |  |
| Clinical Pharmacology | - | 0 | 4 (C) | - |
| Drug Development Science | - | 0 | 4 (C) | - |
| Pharmacology | - | 0 | 2 (C) | - |
| Biopharmaceuticals | - | - | - | ✓ |
| Pharmaceutical Analysis and Quality Control | - | - | - | ✓ |
| Pharmaceutical Technology | ✓ | - | - | - |
| Pharmacy Practice | ✓ | - | - | - |
| *University College London* |  |  |  |  |
| Medicinal Natural Products and Phytochemistry | - | - | - | ✓ |
| Pharmaceutics | - | 0 | 1 (C) | - |
| Drug Discovery and Development | - | 0 | 2 (C) | - |
| Drug Discovery and Pharma Management | - | 0 | 2 (C) | - |
| Pharmaceutical Formulation and Entrepreneurship | - | 0 | 1 (E) | - |
| Clinical Pharmacy, International Practice and Policy | - | - | - | ✓ |
| Drug Sciences | - | - | - | ✓ |
| ***The University of Tokyo*** |  |  |  |  |
| Pharmaceutical Sciences | - | - | - | ✓ |
| ***Karolinska Institute*** |  |  |  |  |
| Pharmaceutical Medicine | - | 1 (C) | 0 | - |

Dedicated: number of standalone RE courses offered by a program; Integrated: number of courses in a program that integrate RE discussions; N/A: no enough information to assess RE education offerings; C: Core course; E: elective course.

## Discussion

This qualitative study revealed a dearth of research ethics education for master’s level pharmacy programs in Jordan. None of the programs offered a standalone research ethics course. A minority (less than half) of programs offered in Jordan integrated partial research ethics instruction into one of their core courses with a focus on human research ethics and research misconduct. Although these programs discuss research ethics themes in their courses that are aligned with what the NIH require in its mandate, these themes still do not capture the scope of topics required in the NIH guidelines [1] as many were missing and not discussed. The core components of research ethics training as indicated by the 2009 NIH mandates and the U.S. ORI include: mentor-mentee responsibilities, research misconduct, research protections of humans and animals, conflict of interest and bias, ethics of collaborative research, data management, publication and authorship ethics, peer-review, and social responsibilities [1, 13]. Globally, assessing research ethics instruction offerings was more challenging as it was entirely based on information available from online official sources, which most of the time was either incomplete or entirely missing. For global programs that we indicated to offer RE education, it was difficult to identify the topics of RE they discuss based solely on keywords from their courses description. Out of the 20 global programs offered, we were able to assess the RE instruction offerings of only 12 (60%). Two out of the twelve offered a standalone RE core course (2/12, 17%), two did not offer any form of research ethics instruction (2/12, 17%), and eight offered RE education incorporated into one or more courses (8/12, 67%). The ultimate can be further classified into programs that integrate RE education into only one core or elective course (2/12, 17%) or into more than one course (6/12, 50%). Based on these findings, one could conclude that formal RE education tends to be lacking globally as well. However, the lack is greater in Jordan than globally (Table 5).

**Table 5.**
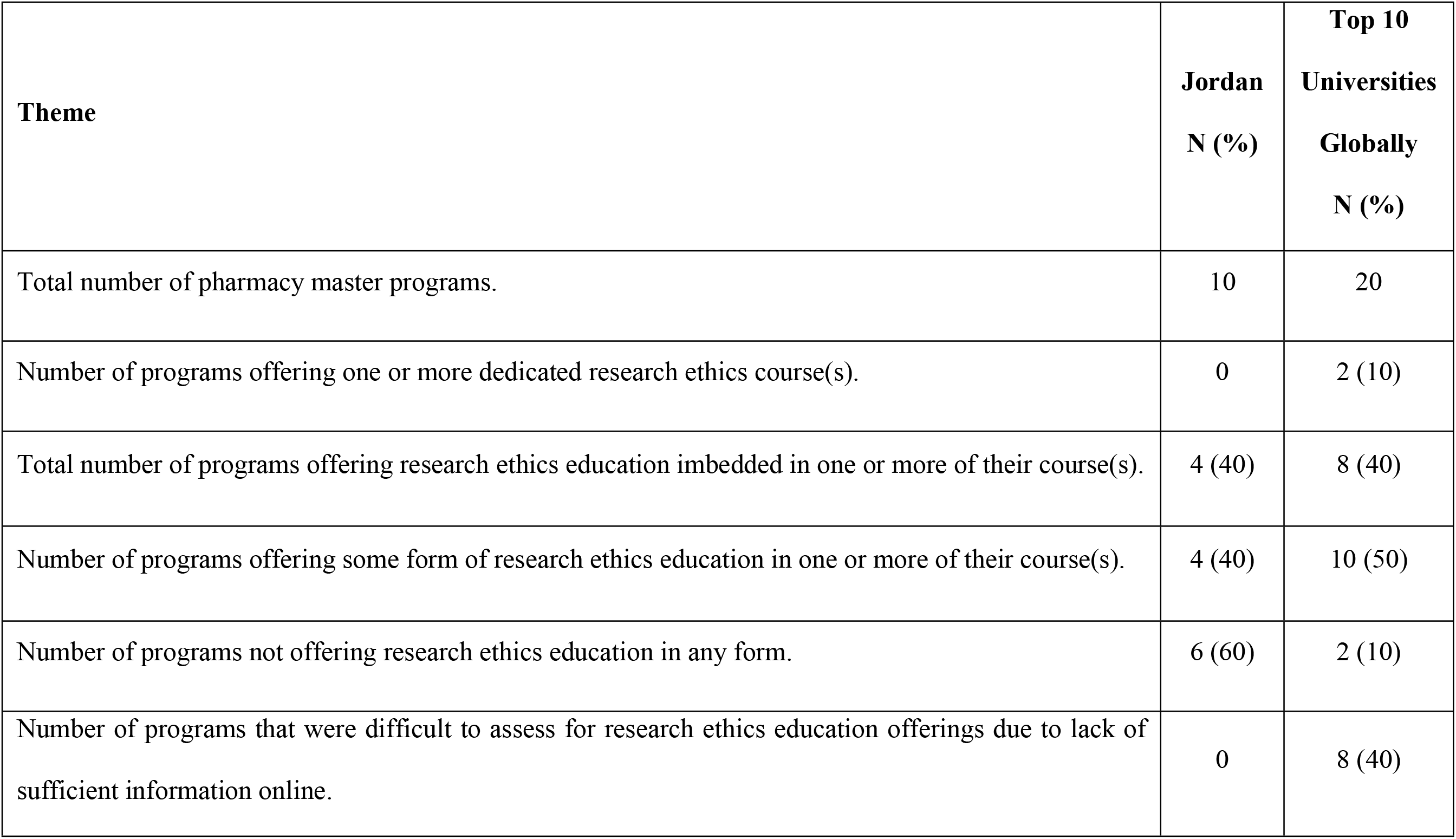
Summary and comparison between research ethics education offerings in Jordan and globally.

Jordan, although considered a developing country, is one of the more academically established countries in the MENA region with progressive research agendas that involve international collaborations. As inter-disciplinary research and collaboration between industrialized and developing countries grows bigger, there emerges the need for capacity building in the developing countries with respect to the responsible conduct of research as well as the ethical review process [23–25]. Numerous deficiencies were previously reported to exist in the ethics guidelines and regulations of countries in low and middle income countries in the Middle East and Africa [26, 27]. A study by Hayder et al. under the former National Bioethics Advisory Commission surveyed health researchers in developing countries to explore issues related to the IRB review. About half of the respondents (44%) disclosed that their projects were not reviewed by an IRB review committee in their countries, of which one-third of these projects were funded by a U.S. funding agency [18].

Systematic education or training in research ethics prior to enrolment in pharmacy graduate education in Jordan is unlikely, as undergraduate pharmacy programs in Jordan do not have a research project completion requirement [28]. Thus, one could assume that pharmacy graduate students lack appropriate training and preparation in the systematic research ethics training most needed at the beginning of the postgraduate program. For that reason, integrating RE education into postgraduate pharmacy programs in Jordan is highly encouraged. In fact, there is a support for this type of training even among health sciences faculty members in Jordan [29]. In this case, pharmacy graduate programs in JUST and JU could serve as a role model in that they contain RE educational material integrated into one or more of their core courses, although integrating a rather “dedicated” RE course into those master programs is highly encouraged, as it was previously reported that RCR programs conducted separately from the standard curricula were more effective than those imbedded into existing modules [30].

Responsible conduct of research education seems to be of great importance in responding to research misconduct and promoting positive attitudes in research. A multi-institutional survey in the U.S., which involved graduate students among the surveyed participants who were enrolled in RCR courses, reported a wide range of plausible outcomes for RCR courses, which had greater impact on knowledge more than fostering skills or attitudes [31]. Several other studies identified favorable outcomes of RCR education in improving knowledge, attitudes, ethical decision making, self-reported behavior, and sense-making skills [5, 32–34] although these improvements might have been described as being modest [5].

It is worth mentioning that despite the NIH training mandates, several studies failed to provide great evidence of its effectiveness [5, 30, 35, 36]. This could be in part due to the fact that research ethics education goals may not be clearly stated nor explicitly specified at the outset of the RCR programs [16]. Therefore, achievable goals need to be clearly articulated at the outset. Several recommendations were addressed in the Delphi consensus panel report regarding the RCR goals and contents and how to adapt the RCR programs to fit the trainees needs [13]. Kalichman and Plemmons have also recommended potential goals for RCR education [16]. Another influencing factors for the effectiveness of RCR educations may include lack of consensus about the contents of RCR education across different institutions and programs [16, 37, 38] leading to high variability of development and implementation of RCR instructions [32, 33, 37]. Furthermore, low level of institutional support [14] and uncoordinated initiatives [3] may also have a negative impact on the outcomes of RCR education. In addition to specifying clear goals for RCR education, several other strategies have been proposed in the literature to overcome the obstacles mitigating the outcomes of RCR training, including competence-based development of research ethics instructions [39, 40], applying research-based narratives assignment [41, 42], careful consideration of instructional designs [35, 43], applying the principles of andragogy [44] and leaning theories [43, 45] to RCR education, improving mentoring strategies of RCR educators [9, 46–49], and using situational factors in real research environment rather than classrooms experience [35, 50, 51], as well as online teaching using internet-based courses [52–55].

### Limitations

There are limitations to this research that can be addressed and mitigated with further research. One major limitation was the lack of available information online for some of the global programs. In addition, identification of research ethics education material for global programs was based solely on keywords we highlighted from the course description to reflect research ethics instruction (Table 1). There is the possibility that these keywords may refer to topics unrelated to research ethics (for instance key words such as issues, problems, regulatory, etc.). As such, our model of data collection and analysis, may lead to an overestimation of research ethics education status rather than an underestimation. For the same reason, little can be inferred about the actual content and the range of RCR topics and core areas covered in the global programs. Moreover, we only included master programs that have mixed course-load and research mandates. Programs that were completed entirely by coursework were excluded as they do not require students to carryout research. Programs entirely focused on research were excluded as well, as there was not clear information available online regarding the structure of these programs. Besides, this study focused on “formal education/training,” whereas informal training may occur within master programs that required students to conduct research. Using ranking systems other than the QS World University Ranking (2017) may lead to results that are slightly different, as the top 10 universities by subject of pharmacy and pharmacology are slightly different depending on the ranking system used.

It is worth mentioning that data collection from local programs in Jordan was more feasible compared to the global programs as it was easier to reach out to schools within Jordan, when needed. For global programs, we did not try to reach out to schools, rather we used information that was available online.

## Conclusions

Training in the RCR for pharmacy graduate students involved in academic research is lacking to higher extent in Jordan than globally as indicated in this qualitative study. Integrating a standalone RCR courses into pharmacy graduate programs and widening the scope of core RCR topics discussed in existing RCR-based courses is highly encouraged. On the other hand, newly established RCR training programs in the MENA region such as the “Research Ethics Program in Jordan; REPJ” which was established in 2015 and is supported by the NIH Fogarty International Center to target young researchers from the MENA region, could play an important role in building capacity among the next generation of scientists. The RCR-Jordan fellows in return would play a pivotal role in raising awareness towards the importance of RCR education and fostering a research culture in which the RCR principles are expected and accepted.

## Acknowledgement

WSA is a participant in the Research Ethics program in Jordan supported by grant # 5R25TW010026-02 from Fogarty International Center of the U.S. National Institute of Health.

## Author contribution

Conceptualization: CN WSA

Formal analysis: WSA

Investigation: WSA CN

Methodology: CN WSA

Project administration: CN

Supervision: CN

Writing – original draft: WSA

Writing – review& editing: CN WSA

## List of abbreviations

RCR: Responsible Conduct of Research
JU: The University of Jordan
JUST: Jordan University of Science and Technology
MENA: Middle East and North Africa
NIH: National Institute of Health
FIC: Fogarty International Center
MSc: Master of Science
U.S.: The United States
QS: Quacquarelli Symonds
ISS: International Student Survey
MERIT: Middle East Research Ethics Training Initiative
NSF: National Science Foundation
NIFA: National Institute of Food and Agriculture
USDA: The United States Department of Agriculture
PHS: Public Health Service
REPJ: Research Ethics Program in Jordan.

## Supporting information

**S1 Table. Research ethics education offerings of pharmacy master programs in Jordan. Listed are universities that offer master programs with mixed coursework and research requirements**.

**S2 Table. Research ethics education offerings of pharmacy master programs in the top 10 universities globally by subject of pharmacy and pharmacology. Listed are all top 10 universities and all pharmacy master programs they offer by mixed coursework and research requirements**.

